# Body mass, temperature, and depth shape the maximum intrinsic rate of population increase in sharks and rays

**DOI:** 10.1101/2021.03.02.433372

**Authors:** Sebastián A. Pardo, Nicholas K. Dulvy

## Abstract

An important challenge in ecology is to understand variation in species’ maximum intrinsic rate of population increase, *r*_*max*_, not least because *r*_*max*_ underpins our understanding of the limits of fishing, recovery potential, and ultimately extinction risk. Across many vertebrates, terrestrial and aquatic, body mass and environmental temperature across important correlates *r*_*max*_ across species. In sharks and rays, specifically, *r*_*max*_ is known be lower in larger species, but also in deep-sea ones. We use an information-theoretic approach that accounts for phylogenetic relatedness to evaluate the relative importance of body mass, temperature and depth on *r*_*max*_. We show that both temperature and depth have separate effects on shark and ray *r*_*max*_ estimates, such that species living in deeper waters have lower *r*_*max*_. Furthermore, temperature also correlates with changes in the mass scaling coefficient, suggesting that as body size increases, decreases in *r*_*max*_ are much steeper for species in warmer waters. These findings suggest that there (as-yet understood) depth-related processes that limit the maximum rate at which populations can grow in deep sea sharks and rays. While the deep ocean is associated with colder temperatures, other factors that are independent of temperature, such as food availability and physiological constraints, may influence the low *r*_*max*_ observed in deep sea sharks and rays. Our study lays the foundation for predicting the intrinsic limit of fishing, recovery potential, and extinction risk species based on easily accessible environmental information such as temperature and depth, particularly for data-poor species.

## Introduction

Ecologists often have to inform policy decisions with knowledge and experience rather than data. A classic and ongoing challenge is that of making a decision about whether a species overfished or not. Overexploitation is the leading cause of extinction risk in the ocean and a major cause also on land (Maxwell *et al*. 2016; Reynolds *et al*. 2005). A large body of work shows that maximum size, either length or weight is a simple heruristic that can guide our thinking about extinction risk. Setting aside the desirability or catchability of different sized fishes, broadly speaking, the larger-bodied species have slower life histories and tend to decline faster when fished (Jennings *et al*. 1998; Reynolds *et al*. 2005). Body size data are among the most widely available traits but it is only one dimension of the life histories of indeterminate-growing species (Chichorro *et al*. 2019). The second dimension of life histories tends to be comprised of time-related “speed of life” traits, such as age at maturity, maximum age, and von Bertalanffy growth rates (Juan-Jordá *et al*. 2013). In the few cases where high quality data can be found, it is increasingly clear that speed of life traits are the better correlate of population dynamics and extinction risk (Anderson *et al*. 2011; Juan-Jordá *et al*. 2015).

The maximum intrinsic rate of population increase (of species at small population sizes) *r*_*max*_ is the integration of speed of life traits and “is perhaps the most fundamental parameter in population biology” (Myers & Worm 2005). Estimates of *r*_*max*_ are commonly used limit reference points as they are equivalent to the fishing mortality that will drive a species to extinction as well as defining the maximum rate of population recovery (Hutchings *et al*. 2012; Dulvy *et al*. 2004; Myers *et al*. 1997). Ever larger databases of life history traits make it easier to calculate *r*_*max*_, providing an opportunity to seek ecological explanations for variation in maximum intrinsic rate of population increase. Indeed, Southwood (1977) viewed habitat quality as the template shaping population growth rates. In the ocean, thermal habitat quality changes profoundly with temperature, such that tropical species have faster life histories and dynamics (Munch & Salinas 2009; Juan-Jordá *et al*. 2013) but attain smaller sizes (Cheung *et al.* 2013; Fisher *et al*. 2010; Jennings *et al*. 2008). From a metabolic perspective alone, at high temperature, fishes are squeezed between low oxygen solubility and high oxygen demand due to temperature-forced metabolic rates (Pörtner *et al*. 2017; Rubalcaba *et al*. 2020; Pauly 2021). Furthermore, within a given thermal habitat, aquatic species exhibit a range of oxygen demands as a result of differing lifestyles, activity patterns and trophic levels, which may have consequences for life histories and population dynamics (Killen *et al*. 2010; 2016; Wong *et al*. 2020). Similar patterns have been observed on land, where allometric variation in mammal production rates (analogous to *r*_*max*_) can also be shaped by differences in lifestyle and trophic level (Sibly & Brown 2007, note that temperature is not an axis of variation in endotherms).

The sharks, rays, and chimaeras (class Chondrichthyes, hereafter referred to as “sharks and rays”) are an ideal taxon for studying the aquatic allometry of demographic rates as they encompass a broad range of sizes and occur across a wide range of temperatures and habitats. The lack of a pelagic larval stage in sharks and rays allows for a more straightforward estimation of population productivity with limited life-history data than their bony counterparts (Myers & Mertz 1998; Pardo *et al*. 2016). Maximum intrinsic rate or population increase *r*_*max*_ among sharks and rays is known to decrease with increasing size (Hutchings *et al*. 2012; Dulvy *et al*. 2014b) and depths (García *et al*. 2008; Simpfendorfer & Kyne 2009). However, previous work has only considered categorical habitat classifications, and there is some evidence that the relationship between *r*_*max*_ and body size breaks down in the deep sea as even small deep-water sharks and rays have very low *r*_*max*_ (Forrest & Walters 2009; Rigby & Simpfendorfer 2015), *suggesting there are stronger constraints on the mass scaling of r*_*max*_ occurring at greater depths than those imposed by body mass and temperature. Thus, depth provides an interesting environmental gradient of energy availability (among other things) to explore interspecific variation in *r*_*max*_.

Here, we examine the separate roles of temperature and depth on the maximum intrinsic rate of population increase and mass scaling relationship among sharks and rays by using estimates of temperature-at-depth derived from species distributions maps (thermal habitat templates) with an information-theoretic approach, while accounting for phylogenetic non-independence.

## Methods

### Data

To determine the role of temperature, depth, and the scaling of body mass on maximum intrinsic rate of population increase, *r*_*max*_ of marine sharks and rays, we gathered three types of data: life-history parameters (maximum weight and *r*_*max*_), species-specific environmental traits (median depth and temperature of occurrence), and phylogenetic trees to account for evolutionary relationships in the model-fitting process. Maximum reported body mass (in grams) was obtained from FishBase (Froese & Pauly 2016) using the rfishbase package (Boettiger *et al*. 2012). We used maximum weight instead of weight at maturity as it is more readily available in the literature, and at a first approximation they are proportional to each other. When body mass data were unavailable, maximum length data were converted to body mass using species-specific length-to-weight conversions also sourced from FishBase. Data for *r*_*max*_ were obtained from a modified Euler-Lotka model following Pardo *et al*. (2016).

Median depth estimates for each species as reported from Dulvy *et al*. (2014a), and temperature-at-depth were derived from species distributions maps. These maps were obtained from AquaMaps which is an online resource of global species distribution models for over 25,000 aquatic species (Kaschner *et al*. 2015).

The core distributions for each species (i.e. where probability of occurrence >= 0.9) were overlaid with the International Pacific Research Center’s interpolated dataset of gridded mean annual ocean temperatures across 27 depth levels (0-2000 m below sea level), which is based on measurements from the Argo Project (see http://apdrc.soest.hawaii.edu/projects/Argo/data/statistics/On_standard_levels/Ensemble_mean/1x1/m00/index.html for data and more information). This temperature interpolation covers most of the world’s oceans, but has incomplete coverage of some shallow coastal areas, such as the Indo-Pacific Triangle and the southeast coast of South America.

To calculate a temperature that characterizes the thermal distribution of each species, we selected the depth level from the grid that was closest to the species’ median depth, and from that grid extracted all the temperature grid points that overlaid the species’ core distribution. From this distribution of temperature values, we calculated the median and set as the temperature value for each species. For example, if a species’ median depth was 130 m we used temperatures from the 150 m depth layer to estimate median annual temperature as 130 m is closer to 150 than 100 m. In species that are known to be mesotherms (e.g. family Lamnidae) we added a correction factor of 3.5 °C.

The phylogenetic trees were obtained from Stein *et al*. (Stein *et al*. 2018) and we followed their scientific nomenclature. Their analyzes did not provide a single tree but a distribution of possible trees with the same topology but differing branch lengths. We ran our analyses by using 20 different trees sampled from their distribution. The results were almost identical regardless of which tree was used, and therefore we only report our findings when using only a single tree (see Table S1 in the Supplementary materials).

### Metabolic scaling expectations

The availability of consistent estimates of environmental temperature derived from species distributions maps allow us to draw from metabolic scaling theory. Savage *et al*. (2004, see equation 8) showed that across multiple taxa, *r*_*max*_ is related with body mass and temperature,

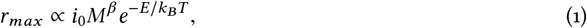

where *r*_*max*_ is the maximum intrinsic rate of population increase (in year^-1^), *i*_0_ is a taxon-specific normalization constant, *M* is the adult body mass of a species (in grams), *β* is the scaling exponent, *E* is the activation energy, *k*_*B*_ is the Boltzmann constant (8.617 × 10^−5^ eV), and *T* is temperature (in Kelvin). The equation above can be simplified when transformed to log-space:

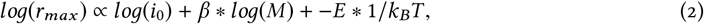

which is equivalent to a simple linear model

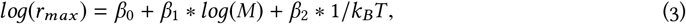

where the intercept *β*_0_ is the log-transformed normalization constant, the coefficient *β*_1_ is the mass scaling, and *β*_2_ is the negative activation energy −*E*, which is the coefficient of inverse temperature 1/*k*_*B*_*T*. We do not use a taxon-specific normalization constant (*i*_0_ = *β*_0_) as we are using a phylogenetic covariance matrix instead of a taxonomic nested structure. Equation 3 is what would be expected from metabolic scaling theory, and is one of the multiple hypotheses we compare in this study.

### What is the role of temperature and depth in the mass scaling relationship of r_max_ ?

We compared nine different models, each representing a different hypothesis of how *r*_*max*_ might vary with body mass, depth, and temperature, using an information-theoretic approach (Burnham & Anderson 2002). We scaled and centered the data for temperature and depth.

To assess whether the estimated median temperature for each species was indeed informative, we also included a model where *r*_*max*_ is explained only by variation in body mass. We compared all models using the corrected Akaike Information Criterion (AICc). If including a parameter improved a model’s AICc value by less than 2, we considered that parameter to be uninformative (Arnold 2010).

To account for non-independence among closely related species, we fitted phylogenetic linear models using the pgls function in the caper package (Orme *et al*. 2013), running on R version 3.3.2 (R Core Team 2016). The phylogenetic covariance matrix, which is contrasted with the residuals from each model, was adjusted using Pagel’s *λ*. This allows for encompassing a wide range of covariance structures corresponding to different trait phylogenetic signal values under a Brownian mode of trait evolution, ranging from no signal at all (i.e. classic OLS; *λ* = 0) to Brownian motion (*λ* = 1) (Revell 2010).

We tested for collinearity by estimating variance-inflation factors (VIF) for all coefficients in the models explored using the car package (Fox & Weisberg 2011). None of the VIF values estimated were greater than 2 (except when interactions are included, which is expected). However, depth and temperature are positively correlated (Pearson’s *r* = 0.64); although this correlation is lower than the threshold of |*r* | > 0.7, where collinearity severely distorts model estimation (Dormann *et al*. 2013), providing further evidence that our models are robust to collinearity.

## Results

We collected temperature, body mass, depth, and *r*_*max*_ data for 63 chondrichthyan species including 40 sharks, 20 rays, and three chimaeras (Fig. 1).

**Figure 1:**
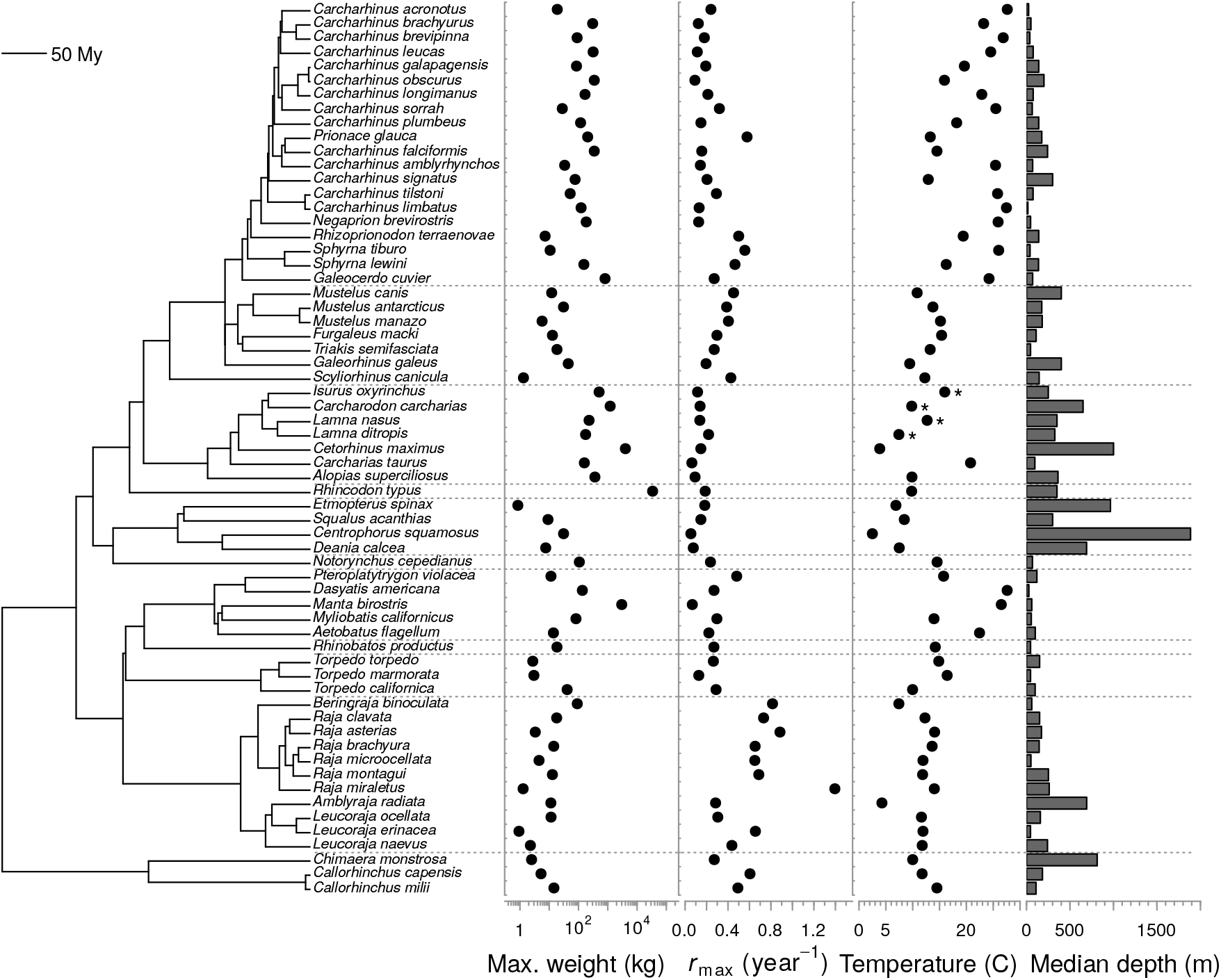
Phylogeny, body mass, maximum intrinsic rate of population increase (*r*_*max*_), temperature, and median depth in sharks, rays, and chimaeras. Phylogenetic tree is based on from Stein *et al*. (2018), body mass estimates were sourced from FishBase (Froese & Pauly 2016), *r*_*max*_ estimates from from Pardo *et al*. (2016), median depth values from Dulvy *et al*. (2014a), and mean annual temperature values (at median depth) were estimated based on species distribution maps from AquaMaps (Kaschner *et al*. 2015) and global temperature grids from the Argo database. Asterisks (*) denote temperature values corrected for mesothermy. Horizontal dotted lines indicate separate taxonomic orders.

The model with the greatest support included temperature and depth as well as an interaction term between body mass and temperature (Table 2). This model also explained the greatest amount of variation (adjusted *R*^2^ = 0.33).

**Table 1:** Models of correlates of maximum intrinsic rate of population increase (*r*_*max*_) tested in our analysis and the hypotheses associated with each.

| Model | Hypothesis |
| --- | --- |
| $\log(r_{max}) \sim \log(M)$ | $r_{max}$ only varies with body mass |
| $\log(r_{max}) \sim \log(M) + depth$ | $r_{max}$ varies with body mass and depth |
| $\log(r_{max}) \sim \log(M) + 1/k_B T$ | $r_{max}$ varies with body mass and temperature |
| $\log(r_{max}) \sim \log(M) + 1/k_B T + depth$ | $r_{max}$ varies with body mass, temperature, and depth |
| $\log(r_{max}) \sim \log(M) + 1/k_B T * depth$ | $r_{max}$ varies with body mass, temperature, and depth, and the effect of temperature varies with depth |
| $\log(r_{max}) \sim \log(M) * depth$ | $r_{max}$ varies with body mass and depth, and the effect of mass (i.e. scaling coefficient) varies with depth |
| $\log(r_{max}) \sim \log(M) * 1/k_B T$ | $r_{max}$ varies with body mass and temperature, and the effect of mass (i.e. scaling coefficient) varies with temperature |
| $\log(r_{max}) \sim \log(M) * depth + 1/k_B T$ | $r_{max}$ varies with body mass, temperature, and depth, and the effect of mass (i.e. scaling coefficient) varies with depth |
| $\log(r_{max}) \sim \log(M) * 1/k_B T + depth$ | $r_{max}$ varies with body mass, temperature, and depth, and the effect of mass (i.e. scaling coefficient) varies with temperature |
| $\log(r_{max}) \sim \log(M) * 1/k_B T + \log(M) * depth$ | $r_{max}$ varies with body mass, temperature, and depth, and the effect of mass (i.e. scaling coefficient) varies with temperature as well as depth |

**Table 2:**
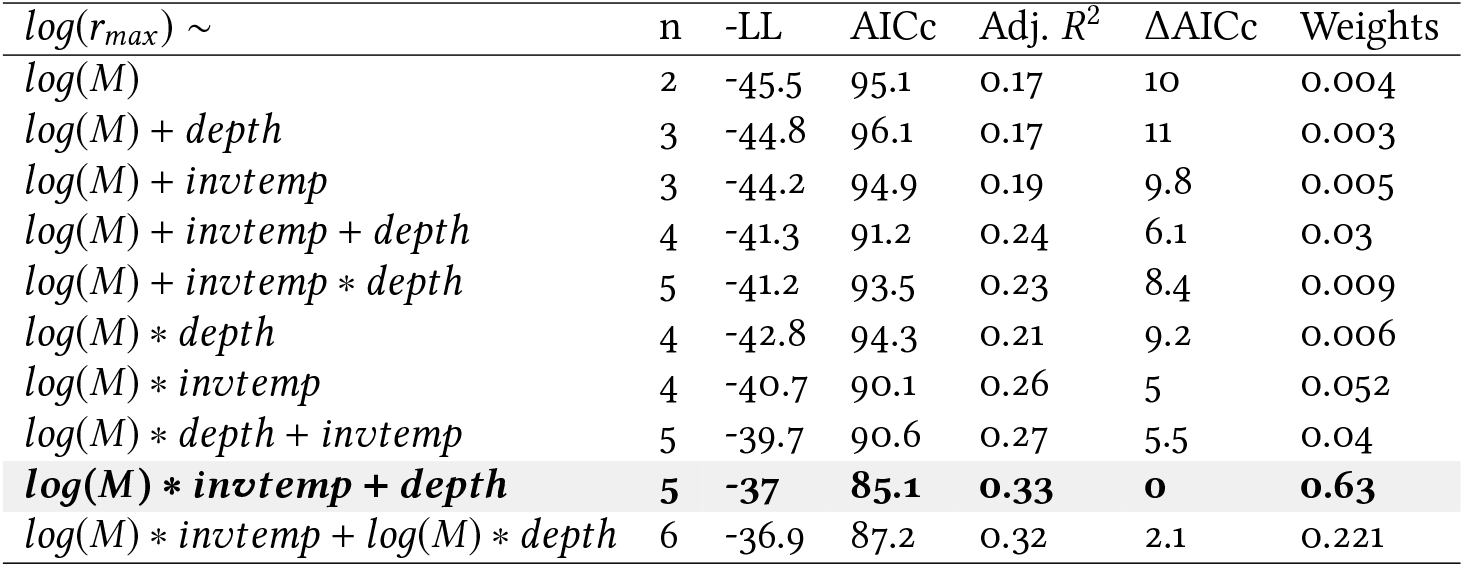
Comparison of *log* (*r*_*max*_) models using standard and corrected Akaike Information Criteria (AICc), number of parameters (n), negative log-likelihood (−LL), adjusted *R*^2^, and Akaike weights. The model with lowest AIC is shown in bold, while the models with ΔAICc ≤ 2 are highlighted in grey.

The model that included the same variables as the top-ranked model as well as an interaction term between body size and depth, received one third the support of the top-ranked model, but had an ΔAICc value greater than 2 without an increase in adjusted *R*^2^. Thus, we considered the interaction between body size and depth to be uninformative and the model was not considered further (Arnold 2010).

All three other models that had marginal support (5 ≤ Δ*AICc* ≤ 7) also included temperature: one was similar to the top-ranked model, but lacked the effect of depth; one included a main effect of temperature, while the effect of body mass varied with depth; and the other includes all the main effects with no interactions (Table 2).

The effect size of body mass in all models is around −0.3 (see shaded areas in Fig. 2, Table 3), which overlaps to the expectation of − ¼ predicted by metabolic scaling theory (Brown *et al*. 2004). Intercepts also consistently overlap zero for all models, albeit with high uncertainty (Table 3). The inclusion of only an additive effect of temperature to the relationship between *r*_*max*_ and body mass was uninformative (Δ*AICc* 10 → 9.8), but inclusion of the interaction between body mass and temperature resulted in an improved model fit (Δ*AICc* = 5.0; Table 2). In contrast, the addition of additive depth alone or an interaction with depth did not improve the relationship between *r*_*max*_ and mass (Δ*AICc* 11 → 9.2); however, the addition of depth to the model with an interaction between body mass and temperature improved this model’s support twelvefold (weight of 0.63 vs. 0.052, Table 2). This effect of depth is consistently negative across the top-ranked models, indicating that *r*_*max*_ decreases with increasing depth.

**Table 3:** Coefficient estimates (95% CI as estimated from standard errors shown in brackets) for all models of *log* (*r*_*max*_). The model with the lowest ΔAICc value is marked in bold and the models with ΔAIC < 2 are highlighted in grey. Pagel’s *λ* is the estimate of branch length transformation on the phylogenetic tree.

| $\log(r_{max}) \sim$ | Intercept | $\log(M)$ | invtemp | depth | $\log(M):invtemp$ | $\log(M):depth$ | invtemp:depth | Pagel's $\lambda$ |
| --- | --- | --- | --- | --- | --- | --- | --- | --- |
| $\log(M)$ | -0.01<br>(-1.01 - 1.00) | -0.31<br>(-0.48 - -0.14) | - | - | - | - | - | 0.85<br>(0.59 - 0.96) |
| $\log(M) + depth$ | -0.03<br>(-1.00 - 0.94) | -0.30<br>(-0.46 - -0.13) | - | -0.19<br>(-0.50 - 0.13) | - | - | - | 0.81<br>(0.46 - 0.96) |
| $\log(M) + invtemp$ | 0.02<br>(-0.99 - 1.03) | -0.33<br>(-0.50 - -0.16) | 0.25<br>(-0.06 - 0.56) | - | - | - | - | 0.87<br>(0.64 - 0.96) |
| $\log(M) + invtemp + depth$ | -0.05<br>(-0.97 - 0.87) | -0.30<br>(-0.46 - -0.15) | 0.49<br>(0.13 - 0.86) | -0.47<br>(-0.84 - -0.11) | - | - | - | 0.80<br>(0.45 - 0.94) |
| $\log(M) + invtemp * depth$ | -0.05<br>(-0.98 - 0.88) | -0.31<br>(-0.47 - -0.15) | 0.52<br>(-0.01 - 1.06) | -0.53<br>(-1.45 - 0.39) | - | - | 0.06<br>(-0.81 - 0.93) | 0.80<br>(0.45 - 0.94) |
| $\log(M) * depth$ | 0.10<br>(-0.85 - 1.04) | -0.31<br>(-0.48 - -0.15) | - | -1.52<br>(-2.87 - -0.18) | - | 0.28<br>(0.00 - 0.56) | - | 0.79<br>(0.43 - 0.95) |
| $\log(M) * invtemp$ | 0.09<br>(-0.86 - 1.04) | -0.33<br>(-0.48 - -0.17) | -1.75<br>(-3.24 - -0.25) | - | 0.40<br>(0.11 - 0.69) | - | - | 0.85<br>(0.60 - 0.96) |
| $\log(M) * depth + invtemp$ | 0.06<br>(-0.85 - 0.97) | -0.32<br>(-0.48 - -0.16) | 0.45<br>(0.09 - 0.81) | -1.54<br>(-2.84 - -0.25) | - | 0.23<br>(-0.04 - 0.51) | - | 0.80<br>(0.45 - 0.94) |
| **$\log(M) * invtemp + depth$** | **0.07**<br>**(-0.79 - 0.93)** | **-0.31**<br>**(-0.46 - -0.16)** | **-1.55**<br>**(-2.98 - -0.12)** | **-0.48**<br>**(-0.82 - -0.14)** | **0.41**<br>**(0.13 - 0.69)** | - | - | **0.78**<br>**(0.47 - 0.93)** |
| $\log(M) * invtemp + \log(M) * depth$ | 0.04<br>(-0.83 - 0.91) | -0.31<br>(-0.46 - -0.15) | -1.97<br>(-4.02 - 0.08) | 0.03<br>(-1.78 - 1.83) | 0.50<br>(0.08 - 0.92) | -0.11<br>(-0.50 - 0.28) | - | 0.78<br>(0.47 - 0.93) |

**Figure 2:**
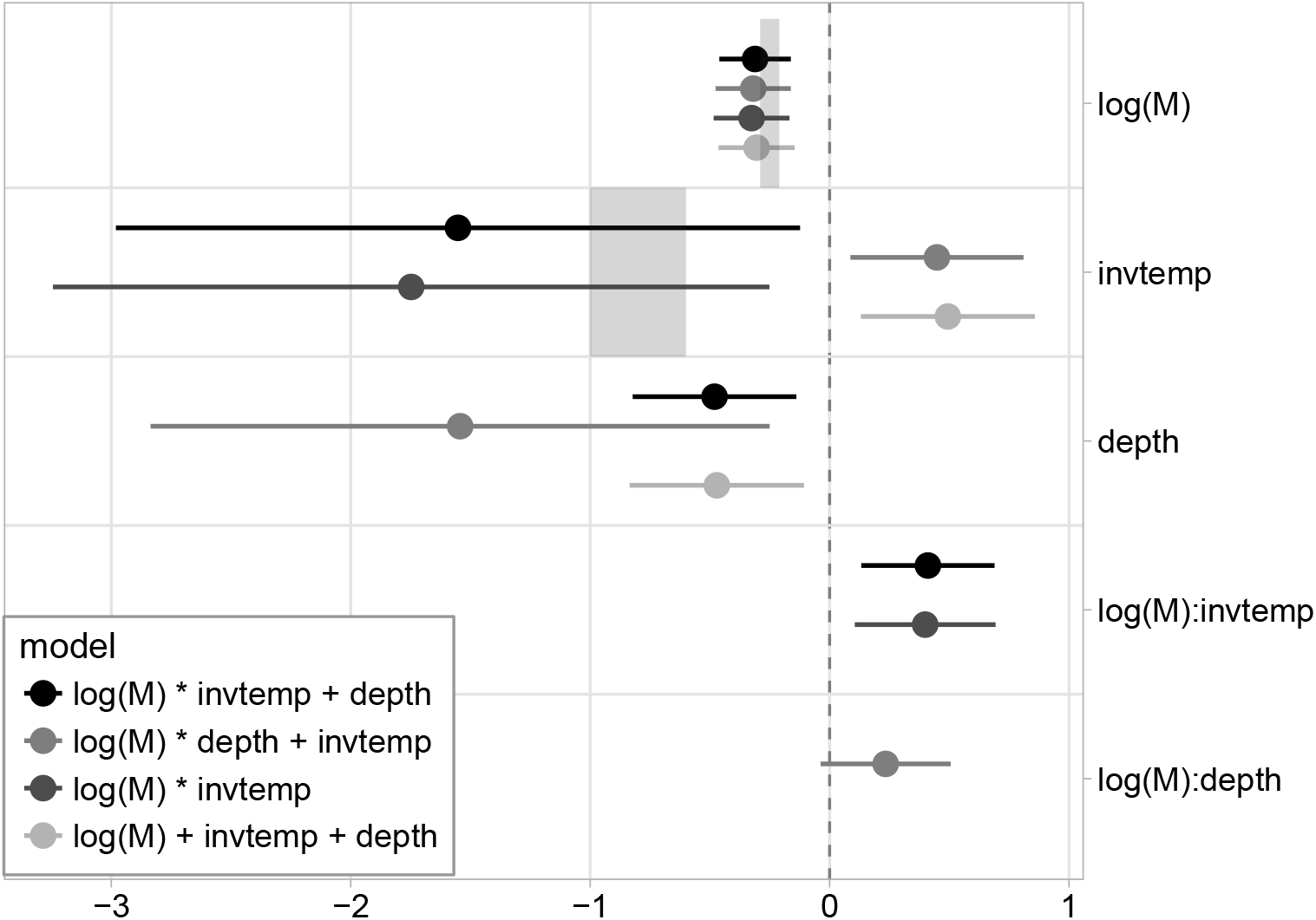
Coefficient plots for the four models of *log* (*r*_*max*_) with lowest AICc values. Lighter colours indicate models with decreasing support based on ΔAICc. Error bars show the 95% confidence intervals, and effect sizes were considered significant when confidence intervals do not overlap zero. Shaded areas show the expected effect sizes for body mass (-0.33 to -0.25) and temperature (-1.0 to -0.6) based on metabolic scaling theory.

The interaction between body mass and temperature was always positive (Table 3) indicating that the mass-scaling coefficient becomes shallower (i.e. slope increasing from a negative value towards zero) with decreasing temperatures. In the top-ranked model, this change in slope with body mass results in the effect of temperature being positive for species up to ∼ 3 kg maximum weight, however for larger species it switches directions and the relationship between temperature on *r*_*max*_ becomes increasingly negative with increasing maximum weight (Fig. 3b). Given that this best model did not have any interaction terms including depth, predicted *r*_*max*_ values decrease with increasing depth (Fig. 3a). Therefore, this model only resulted in plausible temperature effect sizes, based on the expectation from metabolic scaling theory where this coefficient of the inverse of temperature should be negative (−*E* in eq. 2), for species less than 3 kg in size. However, this interpretation of the coefficient direction needs to be taken with caution given that including interactions might be more difficult to translate into a mechanistic understanding of biological rates.

**Figure 3:**
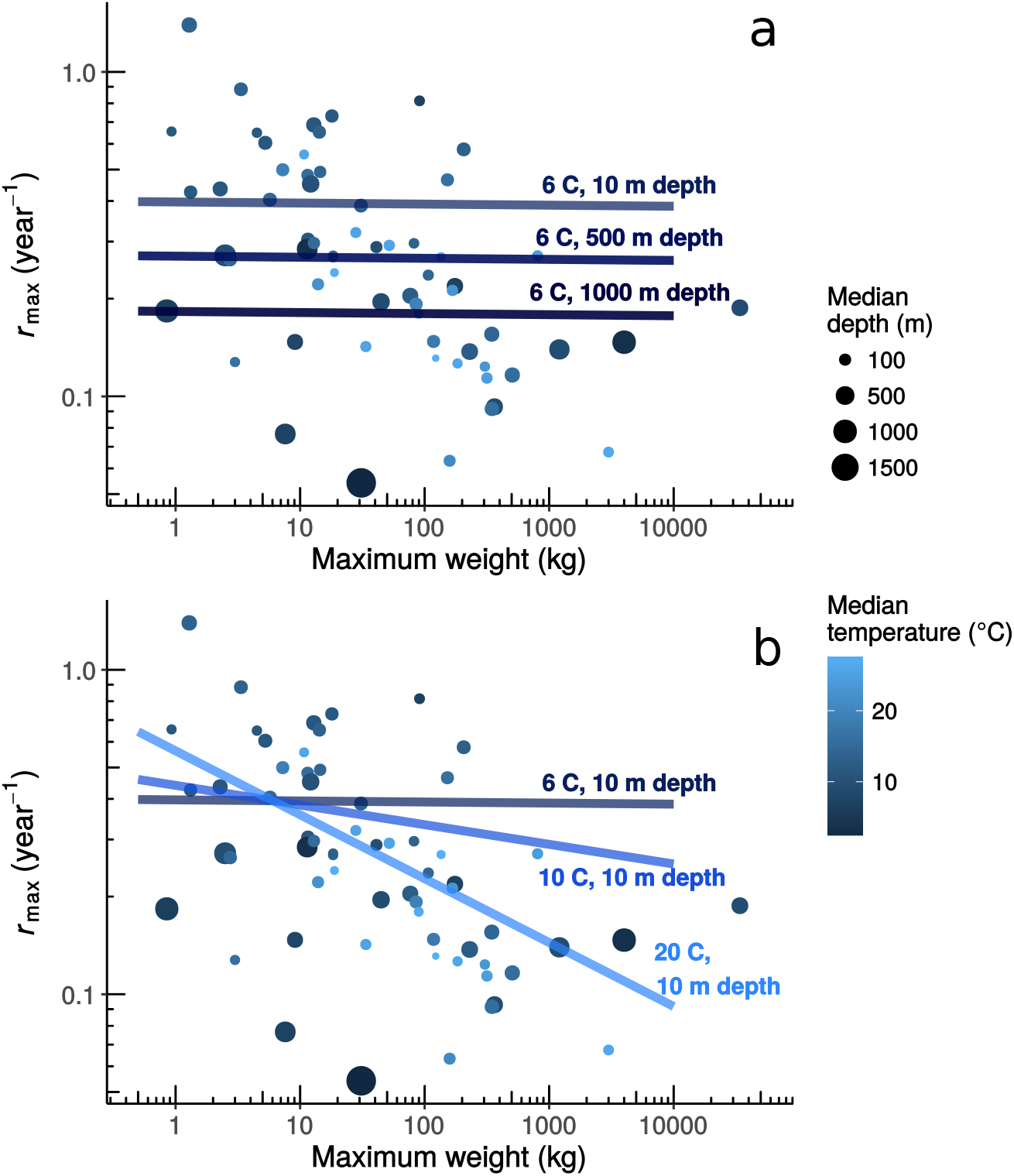
Relationship between maximum weight and maximum intrinsic rate of population increase *r*_*max*_, in log space, for 63 chondrichthyan species. Median depth and temperature for each species are shown by the point size and colour, respectively. Median temperatures are corrected for species which have body temperatures that are higher than their surroundings. Fitted lines show predicted relationships based on the top-ranked model. (a) Predicted allometric changes of *r*_*max*_ across median depths (10, 500, 1000 m) but constant median temperature (6 C), and (b) predicted allometric changes of *r*_*max*_ for three different median temperatures (6, 10, 20 C) but constant median depth (10 m).

There was a strong phylogenetic signal from the residuals of *r*_*max*_ in all ten models examined. Estimated *λ* ranged between 0.78 and 0.87 with the best model having the lowest *λ* value (Table 3).

## Discussion

We generally find that maximum intrinsic rate of population increase is lower in larger bodied species. However, we also found two intriguing depth and temperature related effects. First, we found that *r*_*max*_ decreases at greater depths across marine sharks and rays. Further, the relationship between mass and *r*_*max*_ is temperature-dependent, such that *r*_*max*_ decreases with increasing body mass more steeply at higher temperatures. We also found that the scaling of body mass and *r*_*max*_ changes across a temperature gradient, which is in line with the observation by Rigby & Simpfendorfer (2015) and suggests the absence of a relationship between size and *r*_*max*_ in deep-sea sharks and rays is perhaps due to the low temperatures found in the deep sea rather than depth itself. In other words, it is temperature rather than depth that “flattens” the mass-scaling of *r*_*max*_ in sharks and rays, and species that live in colder environments have a shallower mass-scaling relationship with *r*_*max*_ than those that live in warmer waters. Overall, the interaction between the effect of body mass and temperature likely drives the non-linear scaling of shark and ray *r*_*max*_. Our findings imply that body size might not be a good indicator of *r*_*max*_ among cold-water shark and ray species, which is an important consideration for the conservation and management of these often data-poor species.

There is increasing evidence that the allometries of biological rates can be non-linear, which can stem from multiple separate mechanisms acting concurrently (Kolokotrones *et al*. 2010; Glazier 2005; 2015). Thus, differences in temperature could potentially affect scaling relationships via separate mechanisms (Ohlberger *et al*. 2012; Bruno *et al*. 2015). The differing effects of physiological, energetic, and geometric processes on biological rates across allometries could explain the non-linearity of the relationship observed in this study and are discussed below.

As dissolved oxygen concentrations decrease as water temperature increases (Pauly 2010), the largest bodied species and individuals may become more oxygen limited, which potentially explains why *r*_*max*_ decreases more steeply with increasing body size in warmer water species than colder water ones (Rubalcaba *et al. 2020*). One mechanism that might plausibly account for this is the gill-oxygen limitation theory (GOLT, Pauly & Cheung 2018; Pauly 2021). According to this theory, gill surface area imposes a stronger limitation on oxygen uptake as organisms grow larger, which results in less energy allocated to growth and reproduction under warmer conditions (i.e. waters with less dissolved oxygen). While physiologists dispute the directionality of this mechanism (e.g. Lefevre *et al*. 2018; *Seibel & Deutsch 2020*) it is increasingly clear that individuals with ablated gills have lower metabolic rate and maximal metabolic rate may be depressed in larger individuals in warmer (Pauly 2010; Rubalcaba *et al*. 2020). Assuming that (gill) oxygen limitation is stronger in larger species than smaller ones, oxygen uptake might not be limiting in smaller species where other factors might be shaping *r*_*max*_. Thus, the positive effect of temperature on *r*_*max*_ in smaller species might be a direct result of increased speed of chemical and metabolic reactions as hypothesized by metabolic theory (Brown *et al*. 2004; Savage *et al*. 2004). In larger species, the positive effect of temperature on metabolic reactions could be completely overridden by the oxygen limitation that gills pose as organisms increase in volume (Pauly & Cheung 2018), resulting in a negative relationship between temperature and *r*_*max*_. Nonetheless, some species might be behaviourally evading these temperature (and oxygen) constraints. For example, the Whale Shark (*Rhincodon typus*) is the largest extant fish and lives in warm waters throughout the world’s oceans, yet it performs deep dives (>1000 m) into cold waters which are hypothesized to aid in thermoregulation to dissipate excess heat (Thums *et al*. 2012), and our finding leads us to speculate that they perhaps also access waters with higher oxygen concentrations. While the effect of oxygen concentrations across depth on *r*_*max*_ was not explored in this manuscript, it might further explain the low *r*_*max*_ estimates seen among deep sea species in line with the expectations from the GOLT and other general theories of oxygen limitation (Pörtner *et al*. 2017; Deutsch *et al*. 2020; Rubalcaba *et al*. 2020; *Pauly 2021*).

The existence of a depth dimension to *r*_*max*_, albeit complex, is worth exploring further. Many potential mechanisms could drive the decrease in *r*_*max*_ among chondrichthyans with increasing depth, all of which relate to the amount of energy available for metabolic processes: temperature, light, and consequently primary production decrease below the photic zone (Gage & Tyler 1991; Jahnke 1996); metabolic capacity and requirements decrease with depth as dissolved oxygen and activity levels decrease (Childress *et al*. 1990; *Seibel & Drazen 2007*); animal biomass, and hence food availability, also decreases with increasing depth (Rex *et al*. 2006); even the unique physiology of sharks and rays can increase the energetic cost of living in the deep, and consequently reduce the energy available for production (Treberg & Speers-Roesch 2016). Comparative studies of metabolic rate in marine organisms suggest that the lower metabolic rate seen in deep-water (>1000 m) species is a result of the “visual-interactions hypothesis” (Childress *et al*. 1990; *Seibel & Drazen 2007*), rather than lower food availability or other factors. This hypothesis posits that deep-sea species have lower activity levels as they interact with predators and prey on much smaller spatial scales, or less frequently, than shallow water ones, as the distance at which predators and prey interact is reduced as light levels drop, resulting in lower basal metabolic rates and hence lower productivities (Drazen & Seibel 2007), and also shallower mass-scaling of metabolic rate in deeper waters (Rubalcaba *et al*. 2020). A decrease in *r*_*max*_ with increasing depth could also arise from the unique physiological challenges on chondrichthyans imposed by living in the deep, that limit their ability to use available energy for production. Treberg & Speers-Roesch (2016) hypothesized two separate physiological constraints limiting energy availability in deep water chondrichthyans: the energetic cost of lipid accumulation for buoyancy, and nitrogen limitation due to their osmoregulatory strategy. These physiological limitations could be acting in concert with energetic limitations of low food availability to reduce even further the capability of deep-sea chondrichthyans to grow in size and produce offspring. The complex energetic gradients discussed above can potentially affect the productivity and population growth rates of species found at different depth, even after temperature is accounted for, and as such, depth can be thought of as an added axis of variation in the slow-fast life-history continuum (Sibly & Brown 2007).

The strong phylogenetic signal in the residuals of *r*_*max*_ suggests that other aspects of biology shape their maximum population growth rate, which are likely evolutionarily conserved traits. This likely stems, in part, from estimates of *r*_*max*_ being strongly influenced by reproductive output (Pardo *et al*. 2018), which is strongly conserved among closely related species (Dulvy & Reynolds 1997). This strong phylogenetic signal opens up the door for predictive modelling of *r*_*max*_ based on phylogenetic relationships.

While our study is a first attempt at understanding the relationship between *r*_*max*_, size, and the environment, there is a considerable amount of unexplained variation in our models. Considering the caveats when interpreting the findings of this study might enable future studies to improve model fits. First, there is considerable amount of collinearity between depth and temperature (Pearson’s *r* = 0.64). This is somewhat unavoidable as warmer waters are almost chiefly found in shallow depths, while deep waters are consistently cold. While the diagnostic tests we performed suggest that our results are robust to this collinearity, it is important to bear this in mind, particularly if our framework is to be adapted in the future for predictive purposes. Second, our study is limited by the choice of hypotheses being examined. While we competed nine different hypotheses using an information theoretic approach, it is possible that other hypotheses not considered in this study are better at explaining the patterns in the data. Including other environmental variables mentioned above, such as dissolved oxygen or net primary production, could result in models that are better supported in the data than the models compared in this study. Concomitantly, we only explored simple interactions from a linear modelling framework; other non-linear frameworks for modelling might be better supported by the data. Third, we used maximum weight as our metric of mass as it is the most readily available trait among data-poor sharks and rays, however it is also among the most variable and our choice of this life-history trait might contribute to the low *R*^2^ of our models. Weight at maturity is a likely better parameter to use, albeit more difficult to estimate, as it more precise than estimates of maximum weight, and also aligns better with expectations of metabolic scaling (Savage *et al*. 2004) and life-history theories (Charnov *et al*. 2007). Lastly, our method of accounting for mesothermy (i.e. incrementing environmental temperature by a fixed amount for all mesotherms) might not be the most adequate; there are likely large interspecific diffences in the degree of mesothermy, and also how the temperature differential varies across a range of environmental temperatures.

Our study explored the relationships between body mass, temperature, and depth with the maximum intrinsic rate of increase of sharks and rays and paves the way for further investigation of the mechanisms behind these relationships. For example, investigating how the mass-scaling relationship in chondrichthyans from solely shallow environments varies with temperature might confirm that the apparent “flattening” of the mass scaling relationship of productivity at depth was indeed due to temperature. Taking into account additional spatial datasets that provide information on nutrient availability in the deep sea, such as sedimentation rates or organic carbon burial rates (Jahnke 1996), could provide a better understanding of the degree to which food availability limits productivity among deep-sea sharks and rays. Similarly, incorporating data on dissolved oxygen content would elucidate whether the observed temperature effects, particularly among medium to large-sized species, are a result of oxygen limitation. Alternatively, exploring in more detail whether the effect of depth on *r*_*max*_ levels off after a certain threshold (as it does for basal metabolic rate in pelagic fishes, supporting the visual-interactions hypothesis) or it is continuous would help elucidate the mechanism driving the observed relationship with depth. Identifying the mechanisms behind these observed relationships among body size, temperature, and depth will eventually provide a much clearer understanding of how the productivity, and therefore the vulnerability of species, varies across the world’s oceans.

## Acknowledgements

We are grateful to Jennifer Bigman, Philina English, Sarah Gravel, Daniel Greenberg, Peter Kyne, Florent Mazel, Arne Mooers, and Elan Stopnitzky for their insightful comments on the manuscript and help with the figures. We acknowledge AquaMaps for making their data freely available and especially Kristin Kaschner for discussions on distribution model outputs. SAP & NKD were funded by Natural Sciences and Engineering Research Council, and NKD was also funded by the Canada Research Chairs Program.

## Data accessibility

Data and analyses are available as a GitHub repository:

http://www.github.com/sebpardo/shark-rmax-scaling.

## Notes

https://www.github.com/sebpardo/shark-rmax-scaling

### Competing Interest Statement

The authors have declared no competing interest.

## References

Anderson, S.C., Farmer, R.G., Ferretti, F., Houde, A.L.S. & Hutchings, J.A. (2011). Correlates of vertebrate extinction risk in Canada. BioScience, 61, 538–549.

Arnold, T.W. (2010). Uninformative Parameters and Model Selection Using Akaike’s Information Criterion. The Journal of Wildlife Management, 74, 1175–1178.

Boettiger, C., Temple Lang, D. & Wainwright, P. (2012). rfishbase: exploring, manipulating and visualizing FishBase data from R. Journal of Fish Biology, 81, 2030–2039.

Brown, J.H., Gillooly, J.F., Allen, A.P., Savage, V.M. & West, G.B. (2004). Toward a metabolic theory of ecology. Ecology, 85, 1771–1789.

Bruno, J.F., Carr, L.A. & O’Connor, M.I. (2015). Exploring the role of temperature in the ocean through metabolic scaling. Ecology, 96, 3126–3140.

Burnham, K.P. & Anderson, D.R. (2002). Model Selection and Multimodel Inference: A Practical Information-Theoretic Approach. Second edi edn. Springer New York, New York, NY.

Charnov, E., Warne, R. & Moses, M. (2007). Lifetime Reproductive Effort. The American Naturalist, 170, E129–E142.

Cheung, W.W.L., Sarmiento, J.L., Dunne, J., Frölicher, T.L., Lam, V.W.Y., Deng Palomares, M.L., Watson, R. & Pauly, D. (2013). Shrinking of fishes exacerbates impacts of global ocean changes on marine ecosystems. Nature Climate Change, 3, 254–258.

Chichorro, F., Juslén, A. & Cardoso, P. (2019). A review of the relation between species traits and extinction risk. Biological Conservation, 237, 220–229.

Childress, J., Cowles, D., Favuzzi, J. & Mickel, T. (1990). Metabolic rates of benthic deep-sea decapod crustaceans decline with increasing depth primarily due to the decline in temperature. Deep Sea Research Part A. Oceanographic Research Papers, 37, 929–949.

Deutsch, C., Penn, J.L. & Seibel, B. (2020). Metabolic trait diversity shapes marine biogeography. Nature, 585, 557–562.

Dormann, C.F., Elith, J., Bacher, S., Buchmann, C., Carl, G., Carré, G., Marquéz, J.R.G., Gruber, B., Lafour-cade, B., Leitão, P.J., Münkemüller, T., McClean, C., Osborne, P.E., Reineking, B., Schröder, B., Skidmore, A.K., Zurell, D. & Lautenbach, S. (2013). Collinearity: a review of methods to deal with it and a simulation study evaluating their performance. Ecography, 36, 27–46.

Drazen, J.C. & Seibel, B.A. (2007). Depth-related trends in metabolism of benthic and benthopelagic deep-sea fishes. Limnology and Oceanography, 52, 2306–2316.

Dulvy, N.K., Ellis, J.R., Goodwin, N.B., Grant, A., Reynolds, J.D. & Jennings, S. (2004). Methods of assessing extinction risk in marine fishes. Fish and Fisheries, 5, 255–276.

Dulvy, N.K., Fowler, S.L., Musick, J.A., Cavanagh, R.D., Kyne, P.M., Harrison, L.R., Carlson, J.K., Davidson, L.N., Fordham, S.V., Francis, M.P., Pollock, C.M., Simpfendorfer, C.A., Burgess, G.H., Carpenter, K.E., Compagno, L.J.V., Ebert, D.A., Gibson, C., Heupel, M.R., Livingstone, S.R., Sanciangco, J.C., Stevens, J.D., Valenti, S. & White, W.T. (2014a). Extinction risk and conservation of the world’s sharks and rays. eLife, 3, e00590.

Dulvy, N.K., Pardo, S.A., Simpfendorfer, C.A. & Carlson, J.K. (2014b). Diagnosing the dangerous demography of manta rays using life history theory. PeerJ, 2, e400.

Dulvy, N.K. & Reynolds, J.D. (1997). Evolutionary transitions among egg-laying, live-bearing and maternal inputs in sharks and rays. Proceedings of the Royal Society of London B: Biological Sciences, 264, 1309–1315.

Fisher, J.A.D., Frank, K.T. & Leggett, W.C. (2010). Global variation in marine fish body size and its role in biodiversityfi?ecosystem functioning. Marine Ecology Progress Series, 405, 1–13.

Forrest, R.E. & Walters, C.J. (2009). Estimating thresholds to optimal harvest rate for long-lived, low-fecundity sharks accounting for selectivity and density dependence in recruitment. Canadian Journal of Fisheries and Aquatic Sciences, 66, 2062–2080.

Fox, J. & Weisberg, S. (2011). An R Companion to Applied Regression. 2nd edn. Sage, Thousand Oaks CA.

Froese, R. & Pauly, D. (2016). FishBase.

Gage, J.D. & Tyler, P.A. (1991). Deep-sea biology: a natural history of organisms at the deep-sea floor. Cambridge University Press.

García, V.B., Lucifora, L.O. & Myers, R.A. (2008). The importance of habitat and life history to extinction risk in sharks, skates, rays and chimaeras. Proceedings of the Royal Society B, 275, 83–89.

Glazier, D.S. (2005). Beyond the ‘3/4-power law’: variation in the intra-and interspecific scaling of metabolic rate in animals. Biological reviews of the Cambridge Philosophical Society, 80, 611–62.

Glazier, D.S. (2015). Is metabolic rate a universal fipacemaker’ for biological processes? Biological Reviews, 90, 377–407.

Hutchings, J.A., Myers, R.A., García, V.B., Lucifora, L.O. & Kuparinen, A. (2012). Life-history correlates of extinction risk and recovery potential. Ecological Applications, 22, 1061–1067.

Jahnke, R.A. (1996). The global ocean flux of particulate organic carbon: Areal distribution and magnitude. Global Biogeochemical Cycles, 10, 71–88.

Jennings, S., Mélin, F., Blanchard, J.L., Forster, R.M., Dulvy, N.K. & Wilson, R.W. (2008). Global-scale predictions of community and ecosystem properties from simple ecological theory. Proceedings of the Royal Society B: Biological Sciences, 275, 1375–1383.

Jennings, S., Reynolds, J.D. & Mills, S.C. (1998). Life history correlates of responses to fisheries exploitation. Proceedings of the Royal Society B, 265, 333–339.

Juan-Jordá, M.J., Mosqueira, I., Freire, J. & Dulvy, N.K. (2013). Life history correlates of marine fisheries vulnerability: a review and a test with tunas and mackerel species. In: Marine extinctions - patterns and processes. CIESM Workshop Monograph no. 45 (ed. Briand, F.). CIESM Publisher, Monaco, pp. 113–128.

Juan-Jordá, M.J., Mosqueira, I., Freire, J. & Dulvy, N.K. (2015). Population declines of tuna and relatives depend on their speed of life. Proceedings of the Royal Society B: Biological Sciences, 282.

Kaschner, K., Kesner-Reyes, K., Garilao, C., Rius-Barile, J., Rees, T. & Froese, R. (2015). AquaMaps: Predicted range maps for aquatic species.

Killen, S.S., Atkinson, D. & Glazier, D.S. (2010). The intraspecific scaling of metabolic rate with body mass in fishes depends on lifestyle and temperature. Ecology Letters, 13, 184–193.

Killen, S.S., Glazier, D.S., Rezende, E.L., Clark, T.D., Atkinson, D., Willener, A.S.T. & Halsey, L.G. (2016). Ecological Influences and Morphological Correlates of Resting and Maximal Metabolic Rates across Teleost Fish Species. The American Naturalist, 187, 592–606.

Kolokotrones, T., Savage, V., Deeds, E.J. & Fontana, W. (2010). Curvature in metabolic scaling. Nature, 464, 753–756.

Lefevre, S., McKenzie, D.J. & Nilsson, G.E. (2018). In modelling effects of global warming, invalid assumptions lead to unrealistic projections. Global Change Biology, 24, 553–556.

Maxwell, S.L., Fuller, R.A., Brooks, T.M. & Watson, J.E.M. (2016). Biodiversity: The ravages of guns, nets and bulldozers. Nature, 536, 143–145.

Munch, S.B. & Salinas, S. (2009). Latitudinal variation in lifespan within species is explained by the metabolic theory of ecology. Proceedings of the National Academy of Sciences, 106, 13860–13864.

Myers, R.A. & Mertz, G. (1998). The limits of exploitation: A precautionary approach. Ecological Applications, 8, 165–169.

Myers, R.A., Mertz, G. & Fowlow, P.S. (1997). Maximum population growth rates and recovery times for Atlantic cod, Gadus morhua. Fishery Bulletin, 95, 762–772.

Myers, R.A. & Worm, B. (2005). Extinction, survival or recovery of large predatory fishes. Philosophical Transactions of the Royal Society B, 360, 13–20.

Ohlberger, J., Mehner, T., Staaks, G. & Hölker, F. (2012). Intraspecific temperature dependence of the scaling of metabolic rate with body mass in fishes and its ecological implications. Oikos, 121, 245–251.

Orme, D., Freckleton, R., Thomas, G., Petzoldt, T., Fritz, S., Isaac, N. & Pearse, W. (2013). caper: Comparative Analyses of Phylogenetics and Evolution in R.

Pardo, S.A., Cooper, A.B., Reynolds, J.D. & Dulvy, N.K. (2018). Qyantifying the known unknowns: estimating maximum intrinsic rate of population increase in the face of uncertainty. ICES Journal of Marine Science, 75, 953–963.

Pardo, S.A., Kindsvater, H.K., Reynolds, J.D. & Dulvy, N.K. (2016). Maximum intrinsic rate of population increase in sharks, rays, and chimaeras: the importance of survival to maturity. Canadian Journal of Fisheries and Aquatic Sciences, 73, 1159–1163.

Pauly, D. (2010). Gasping fish and panting squids: oxygen, temperature and the growth of water-breathing animals. vol. 22. Inter-Research, Oldendorf/Luhe, Germany.

Pauly, D. (2021). The gill-oxygen limitation theory (GOLT) and its critics. Science Advances, 7, eabc6050.

Pauly, D. & Cheung, W.W.L. (2018). Sound physiological knowledge and principles in modeling shrinking of fishes under climate change. Global Change Biology, 24, e15–e26.

Pörtner, H.O., Bock, C. & Mark, F.C. (2017). Oxygen-and capacity-limited thermal tolerance: bridging ecology and physiology. The Journal of Experimental Biology, 220, 2685–2696.

R Core Team (2016). R: A Language and Environment for Statistical Computing.

Revell, L.J. (2010). Phylogenetic signal and linear regression on species data. Methods in Ecology and Evolution, 1, 319–329.

Rex, M.A., Etter, R.J., Morris, J.S., Crouse, J., McClain, C.R., Johnson, N.A., Stuart, C.T., Deming, J.W., Thies, R. & Avery, R. (2006). Global bathymetric patterns of standing stock and body size in the deep-sea benthos. Marine Ecology Progress Series, 317, 1–8.

Reynolds, J.D., Dulvy, N.K., Goodwin, N.B. & Hutchings, J.A. (2005). Biology of extinction risk in marine fishes. Proceedings of the Royal Society B, 272, 2337–44.

Rigby, C. & Simpfendorfer, C.A. (2015). Patterns in life history traits of deep-water chondrichthyans. Deep Sea Research Part II: Topical Studies in Oceanography, 115, 30–40.

Rubalcaba, J.G., Verberk, W.C.E.P., Hendriks, A.J., Saris, B. & Woods, H.A. (2020). Oxygen limitation may affect the temperature and size dependence of metabolism in aquatic ectotherms. Proceedings of the National Academy of Sciences, 117, 31963–31968.

Savage, V.M., Gillooly, J.F., Brown, J.H., West, G.B. & Charnov, E.L. (2004). Effects of Body Size and Temperature on Population Growth. The American Naturalist, 163, 429–441.

Seibel, B.A. & Deutsch, C. (2020). Oxygen supply capacity in animals evolves to meet maximum demand at the current oxygen partial pressure regardless of size or temperature. The Journal of Experimental Biology, 223, jeb210492.

Seibel, B.A. & Drazen, J.C. (2007). The rate of metabolism in marine animals: environmental constraints, ecological demands and energetic opportunities. Philosophical Transactions of the Royal Society B: Biological Sciences, 362, 2061–2078.

Sibly, R.M. & Brown, J.H. (2007). Effects of body size and lifestyle on evolution of mammal life histories. Proceedings of the National Academy of Sciences, 104, 17707–17712.

Simpfendorfer, C.A. & Kyne, P.M. (2009). Limited potential to recover from overfishing raises concerns for deep-sea sharks, rays and chimaeras. Environmental Conservation, 36, 97–103.

Southwood, T.R.E. (1977). Habitat, the Templet for Ecological Strategies? Journal of Animal Ecology, 46, 337–365.

Stein, R.W., Mull, C.G., Kuhn, T.S., Aschliman, N.C., Davidson, L.N.K., Joy, J.B., Smith, G.J., Dulvy, N.K. & Mooers, A.O. (2018). Global priorities for conserving the evolutionary history of sharks, rays and chimaeras. Nature Ecology & Evolution, 2, 288–298.

Thums, M., Meekan, M., Stevens, J., Wilson, S. & Polovina, J. (2012). Evidence for behavioural thermoregulation by the world’s largest fish. Journal of The Royal Society Interface, 10, 2012.0477.

Treberg, J.R. & Speers-Roesch, B. (2016). Does the physiology of chondrichthyan fishes constrain their distribution in the deep sea? Journal of Experimental Biology, 219, 615–625.

Wong, S., Bigman, J.S. & Dulvy, N.K. (2020). The metabolic pace of life histories across fishes. bioRxiv, p. 2020.11.16.385559.

