## Supplemental Table 1 for "Body mass, temperature, and depth shape the maximum intrinsic rate of population increase in sharks and rays"

for

These Supplementary Materials include supplementary tables for the publication.

Table S1: Differences in corrected Akaike Information Criteria (ΔAICc) for the models run with 20 different iterations of the phylogenetic tree published in Stein *et al.* (2018). The model with the lowest ΔAICc value in each run is highlighted in grey.

| $log(r_{max}) \sim$ | 1 | 2 | 3 | 4 | 5 | 6 | 7 | 8 | 9 | 10 | 11 | 12 | 13 | 14 | 15 | 16 | 17 | 18 | 19 | 20 |
|---|---|---|---|---|---|---|---|---|---|---|---|---|---|---|---|---|---|---|---|---|
| $log(M)$ | 10.70 | 10.40 | 9.30 | 10.50 | 10.60 | 10.30 | 10.40 | 9.30 | 10.50 | 10.40 | 9.30 | 9.80 | 10.60 | 10.60 | 10.50 | 10.50 | 10.50 | 10.70 | 10.70 | 10.40 |
| $log(M) + depth$ | 11.40 | 11.20 | 10.60 | 11.30 | 11.40 | 11.20 | 11.20 | 10.60 | 11.30 | 11.20 | 10.60 | 11.00 | 11.20 | 11.30 | 11.30 | 11.30 | 11.30 | 11.40 | 11.50 | 11.20 |
| $log(M) + invtemp$ | 10.60 | 10.30 | 9.00 | 10.30 | 10.50 | 10.10 | 10.30 | 9.00 | 10.30 | 10.30 | 9.00 | 9.40 | 10.60 | 10.40 | 10.30 | 10.30 | 10.40 | 10.60 | 10.60 | 10.30 |
| $log(M) + invtemp + depth$ | 6.50 | 6.40 | 6.30 | 6.40 | 6.40 | 6.20 | 6.30 | 6.30 | 6.30 | 6.20 | 6.30 | 6.40 | 6.40 | 6.20 | 6.30 | 6.30 | 6.30 | 6.30 | 6.40 | 6.30 |
| $log(M) + invtemp * depth$ | 8.80 | 8.70 | 8.60 | 8.70 | 8.80 | 8.50 | 8.60 | 8.60 | 8.70 | 8.60 | 8.60 | 8.80 | 8.70 | 8.50 | 8.60 | 8.60 | 8.60 | 8.60 | 8.70 | 8.70 |
| $log(M) * depth$ | 9.40 | 9.10 | 9.00 | 9.30 | 9.30 | 9.20 | 9.30 | 9.00 | 9.30 | 9.20 | 9.00 | 9.30 | 9.00 | 9.30 | 9.30 | 9.30 | 9.20 | 9.20 | 9.40 | 9.20 |
| $log(M) * invtemp$ | 5.50 | 5.30 | 4.20 | 5.30 | 5.40 | 5.30 | 5.30 | 4.20 | 5.30 | 5.40 | 4.20 | 4.30 | 5.50 | 5.60 | 5.30 | 5.30 | 5.40 | 5.60 | 5.50 | 5.20 |
| $log(M) * depth + invtemp$ | 5.60 | 5.40 | 5.80 | 5.50 | 5.50 | 5.40 | 5.50 | 5.80 | 5.50 | 5.40 | 5.80 | 5.80 | 5.40 | 5.50 | 5.40 | 5.40 | 5.40 | 5.30 | 5.60 | 5.50 |
| $\boldsymbol{log(M) * invtemp + depth}$ | 0.00 | 0.00 | 0.00 | 0.00 | 0.00 | 0.00 | 0.00 | 0.00 | 0.00 | 0.00 | 0.00 | 0.00 | 0.00 | 0.00 | 0.00 | 0.00 | 0.00 | 0.00 | 0.00 | 0.00 |
| $log(M) * invtemp + log(M) * depth$ | 2.20 | 2.20 | 1.90 | 2.20 | 2.10 | 2.20 | 2.20 | 2.00 | 2.20 | 2.10 | 1.90 | 2.00 | 2.20 | 2.20 | 2.20 | 2.20 | 2.10 | 2.20 | 2.10 | 2.10 |
